# Synergy between eIF5A and Mg2+ enhances elongation in a defined yeast cell-free translation system with synthetic tRNAs

**DOI:** 10.1101/2025.04.30.651424

**Authors:** Yuanhan Jin

**Affiliations:** School of Medicine, Southeast of University

**Keywords:** synthetic tRNA, eIF5A, Mg2+, Saccharomyces cerevisiae, stop codon readthrough

## Abstract

Cell-free translation platforms enable genetic-code engineering but typically rely on native transfer RNAs (tRNAs). Here we reconstitute Saccharomyces cerevisiae translation using a fully synthetic set of 21 in vitro-transcribed tRNAs—one isoacceptor per canonical amino acid plus the initiator—that collectively decode all 61 sense codons. After individual aminoacylation, this minimal pool supports peptide synthesis in a fully recombinant yeast cell-free translation system at yields comparable to native tRNAs. Translation of long open reading frames stalls without supplementation; adding either eukaryotic initiation factor 5A (eIF5A) or elevated Mg2+ restores activity, and together they increase Nano luciferase (NanoLuc) output by approximately fivefold. Alanine scanning implicates basic residues R27 and R87 in eIF5A, rather than the hypusine side chain, as the principal contributors to P-site tRNA stabilization. Higher Mg2+ alone accelerates elongation but increases UAG/UGA readthrough to ∼15%, revealing a tunable speed–fidelity trade-off with unmodified tRNAs. Installing t6A37 or m1G37 on selected synthetic tRNAs further improves processivity. This minimal, programmable yeast platform enables systematic dissection of tRNA-modification functions and provides a practical route to genetic-code engineering.

## Introduction

Cell-free translation enables mechanism-level dissection of gene expression while avoiding the compositional complexity of lysates. In bacteria, the PURE (protein synthesis using recombinant elements) concept showed that translation can be reconstituted from purified parts and tuned with defined buffers(Shimizu et al., 2001; Carlson et al., 2012; Zemella et al., 2015). Yeast systems have progressed from crude extracts to semi-defined or recombinant formats that better capture eukaryotic elongation and folding(Hodgman & Jewett, 2014; Abe et al., 2020; Nagai et al., 2021), but truly minimal and programmable eukaryotic platforms remain scarce.

A central lever of programmability is the transfer RNA (tRNA) pool. Wobble pairing allows substantial compression, and bacterial studies demonstrated that panels as small as ∼21 in vitro-transcribed (IVT) tRNAs can sustain translation under suitable codon usage(Chin, 2017; Hibi et al., 2020; Katoh & Suga, 2022). Extending such minimality to eukaryotes is harder: decoding rules are stricter and many tRNAs require position-37 modifications (for example, t6A37 and m1G37) to stabilize the codon–anticodon mini-helix and maintain the reading frame(Deutsch et al., 2012; Hou, Masuda & Gamper, 2019; Masuda et al., 2021). Although scalable IVT methods can now produce aminoacylatable yeast tRNAs, it remains unclear whether a 21-tRNA set can decode all 61 sense codons with useful yields in a fully defined Saccharomyces cerevisiae system.

Two variables with pronounced influence on elongation when using minimally modified tRNA pools are eukaryotic initiation factor 5A (eIF5A) and Mg2+. eIF5A, uniquely activated by hypusination, interacts with the translating 80S ribosome and promotes otherwise slow elongation events by stabilizing P-site tRNA(Zanelli et al., 2006; Gregio et al., 2009; Melnikov et al., 2016). Mg2+, in turn, tunes ribosome conformational dynamics and decoding energetics; higher Mg2+ can accelerate elongation yet often increases near-/non-cognate accommodation and stop-codon readthrough, revealing a speed-fidelity trade-off (Yamamoto et al., 2010; Schauss et al., 2021; Yamagami, Sieg & Bevilacqua, 2021). Here, we establish a fully defined yeast cell-free translation platform powered by a minimal set of 21 IVT tRNAs that decode all sense codons. We quantify how eIF5A and Mg2+ shape elongation on long ORFs—eIF5A enhancing processivity while preserving accuracy, elevated Mg2+ boosting yield at a fidelity cost—and identify core eIF5A residues (R27/R87) that stabilize the P-site tRNA. This compact, programmable system enables mechanistic dissection of eukaryotic translation and supports genetic-code engineering.

## Results

### Design and preparation of a minimal IVT-tRNA panel

We assembled a 21-member Saccharomyces cerevisiae IVT-tRNA set—one isoacceptor per canonical amino acid plus the initiator — to cover all 61 sense codons (Fig. 1A). Anticodons were selected to favor accurate wobble (G34/C34 where possible), with three pragmatic choices—Ala(UGC), Pro(UGG), and Ile(UAU)—to maximize aminoacylation and downstream performance; stop codons were left unassigned for readthrough assays. All tRNAs were generated by T7 transcription, minimally processed (5′-leader removal where present), refolded in Mg2+ buffer, and purified to a single band on denaturing PAGE (Fig. 1B). Each species was aminoacylated individually using yeast S100 or recombinant aminoacyl-tRNA synthetases, and charging was quantified by incorporation of radiolabeled amino acids (see Methods). Across the panel, charging efficiencies were comparable to native tRNAs, enabling equimolar pooling for subsequent translation assays (Fig. 1C).

**Figure 1.**
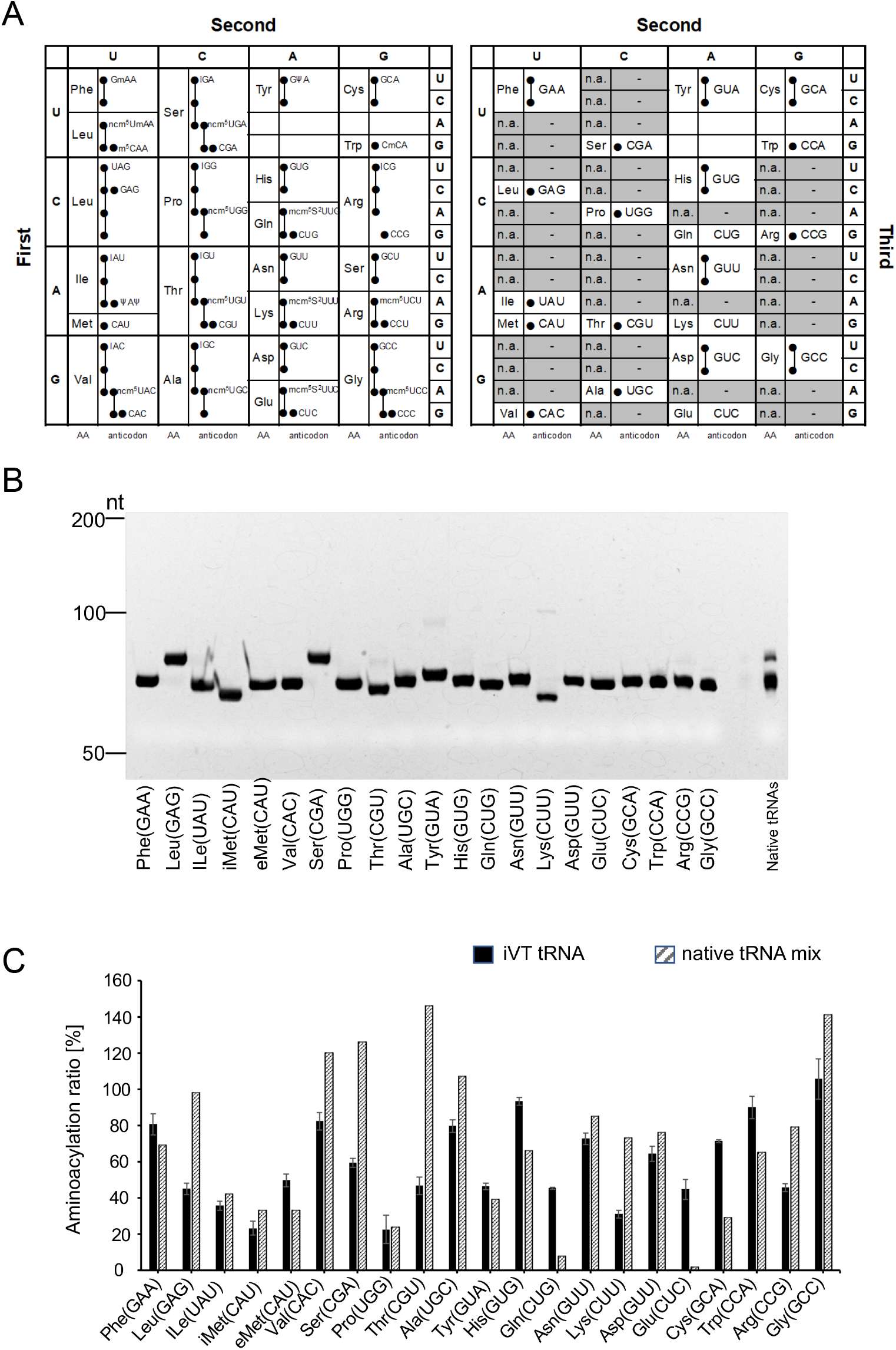
The codon-anti codon selection and constitution of iVT tRNAs. (A) Left: The genetic codes of native yeast tRNAs; Right: selected anticodons for iVT tRNAs and the codons for the mRNA. (B) The Urea-PAGE electrophoresis results of all kinds of puried iVT tRNAs, as well as the native tRNA mix (0.01 A260 units/lane). (C) Amino acid acceptance of each kind of iVT tRNAs under optimal conditions, compared to the respective native tRNA.

### Functional validation of the IVT-tRNA pool in a defined yeast translation system

The pooled IVT-tRNA set supported efficient translation of short reporters, with endpoint signals and kinetics matching reactions assembled with native tRNAs (Fig. 2A). Assay controls confirmed a shared linear range for both pools across time courses and template titrations, with no systematic bias (Supp. Fig. 1A–D).

**Figure 2.**
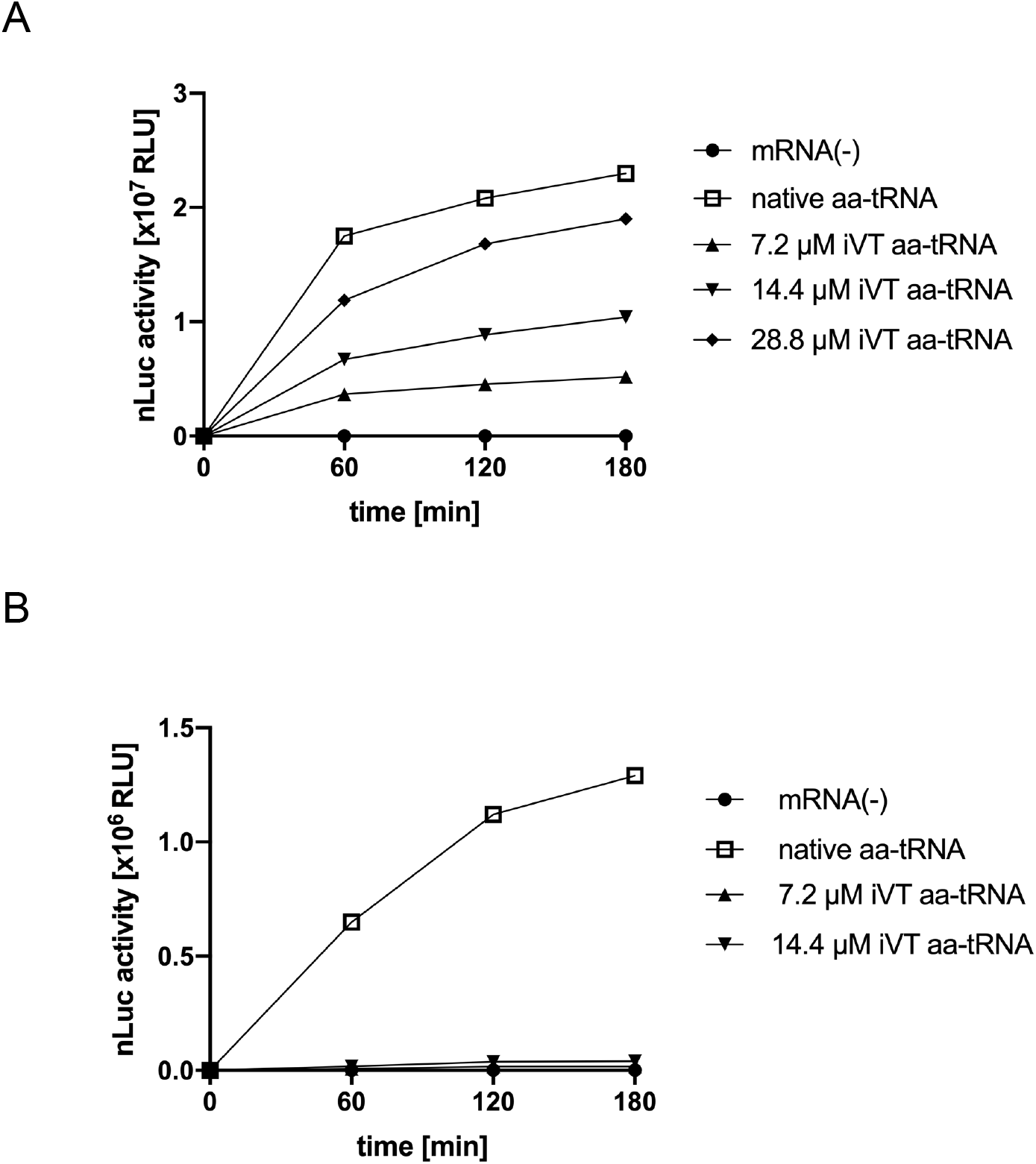
The attempt of translation with iVT tRNAs, compared to the native yeast tRNA mix under 5 mM Mg2+ (without eIF5A). The concentration of native aa-tRNA in the yeast PURE system was 7.2 μM. (A) The synthesis of HiBiT of nanoluciferse. (B) The synthesis of full-length NanoLuc.

By contrast, long ORFs such as NanoLuc (225 aa) produced less full-length product under baseline conditions (Fig. 2B). Supplementing eukaryotic initiation factor 5A (eIF5A) restored elongation: both the hypusinated form [eIF5A(hyp)] and the lysine form lacking hypusine [eIF5A(lys)] increased full-length NanoLuc and reduced truncated intermediates (Fig. 3A–D). Dose–response and time-course series showed that micromolar eIF5A was sufficient, whereas short-ORF outputs changed little—consistent with a primary role in elongation processivity rather than initiation/decoding. Together with the parity controls, these data indicate that the IVT-tRNA pool is interchangeable with native tRNAs for short ORFs, and that eIF5A—independent of hypusination—rescues long-ORF translation in this defined yeast system.

**Figure 3.**
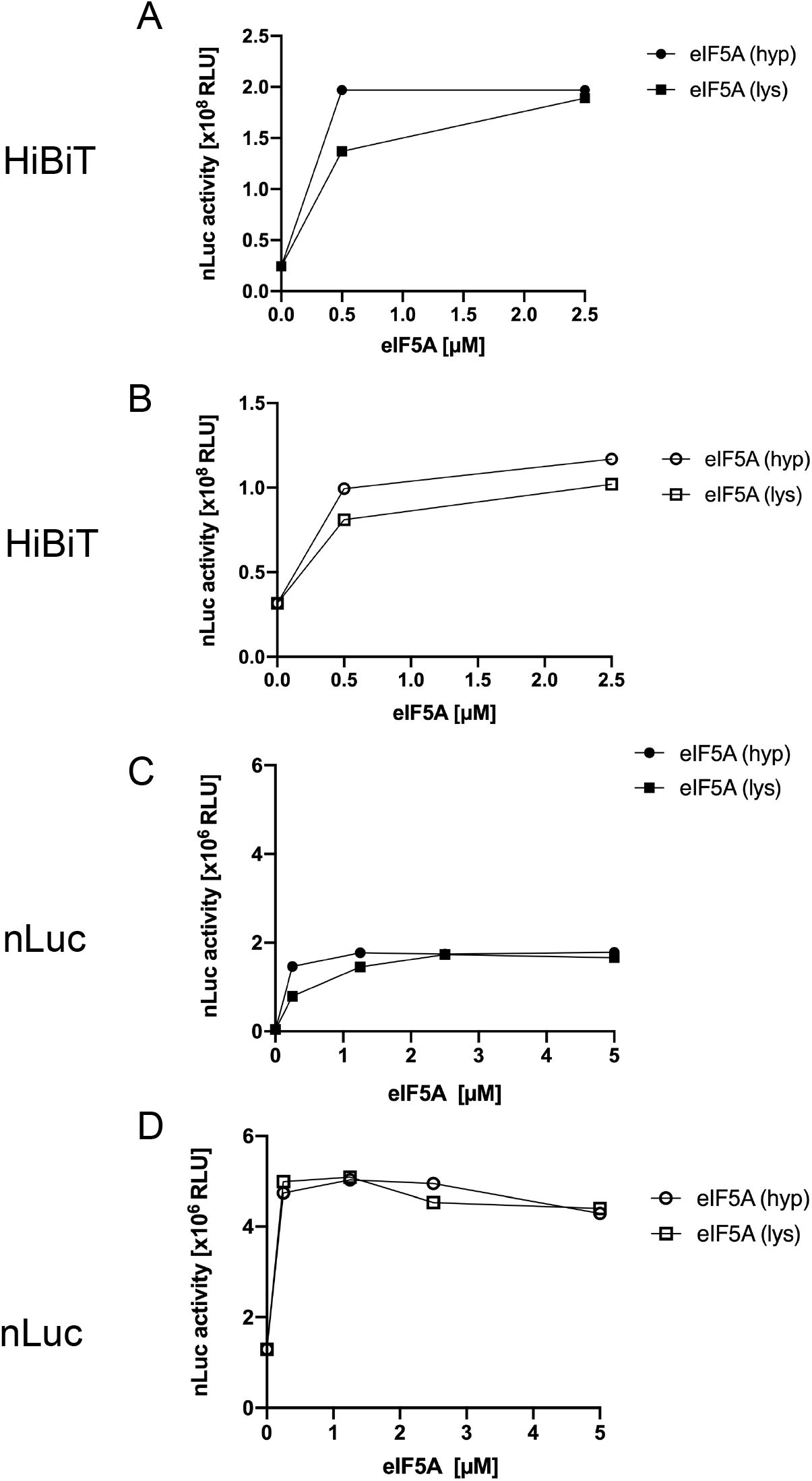
Translation with iVT tRNAs in the presence of eIF5A under 5 mM Mg2+. The activity of NanoLuc was detected after 3hr reaction. (A) The synthesis of HiBiT with iVT aa -tRNAs (14.4 μM) and (B) native aa-tRNA (7.2 μM) in the presence of different concentrations of hypusinated eIF5A [eIF5A (hyp)] or unhypusinated eIF5A [eIF5A (lys)]. (C) The synthesis of full-length NanoLuc with iVT aa -tRNAs and (D) native aa-tRNA in the presence of different concentrations of eIF5A (hyp) or eIF5A (lys).

### Magnesium provides an independent rescue but introduces a fidelity trade-off

Titrating Mg2+ provided an independent route to restore elongation on long ORFs in the absence of eIF5A. Increasing Mg2+ shifted reactions from predominantly truncated products to full-length NanoLuc with a clear dose–response (Fig. 4A), and gel analyses showed a redistribution from stalled/intermediate bands toward the full-length species (Fig. 4B). Time-course comparisons at representative low versus high Mg2+ revealed earlier appearance and faster accumulation of full-length product at higher Mg2+ (Fig. 4C). The rescue generalized across reporter inputs and was observed with both native and IVT tRNA pools (Fig. 4D); replicate titrations and kinetics in Supp. Fig. 2 reproduced these trends and delineated the effective Mg2+ window.

**Figure 4.**
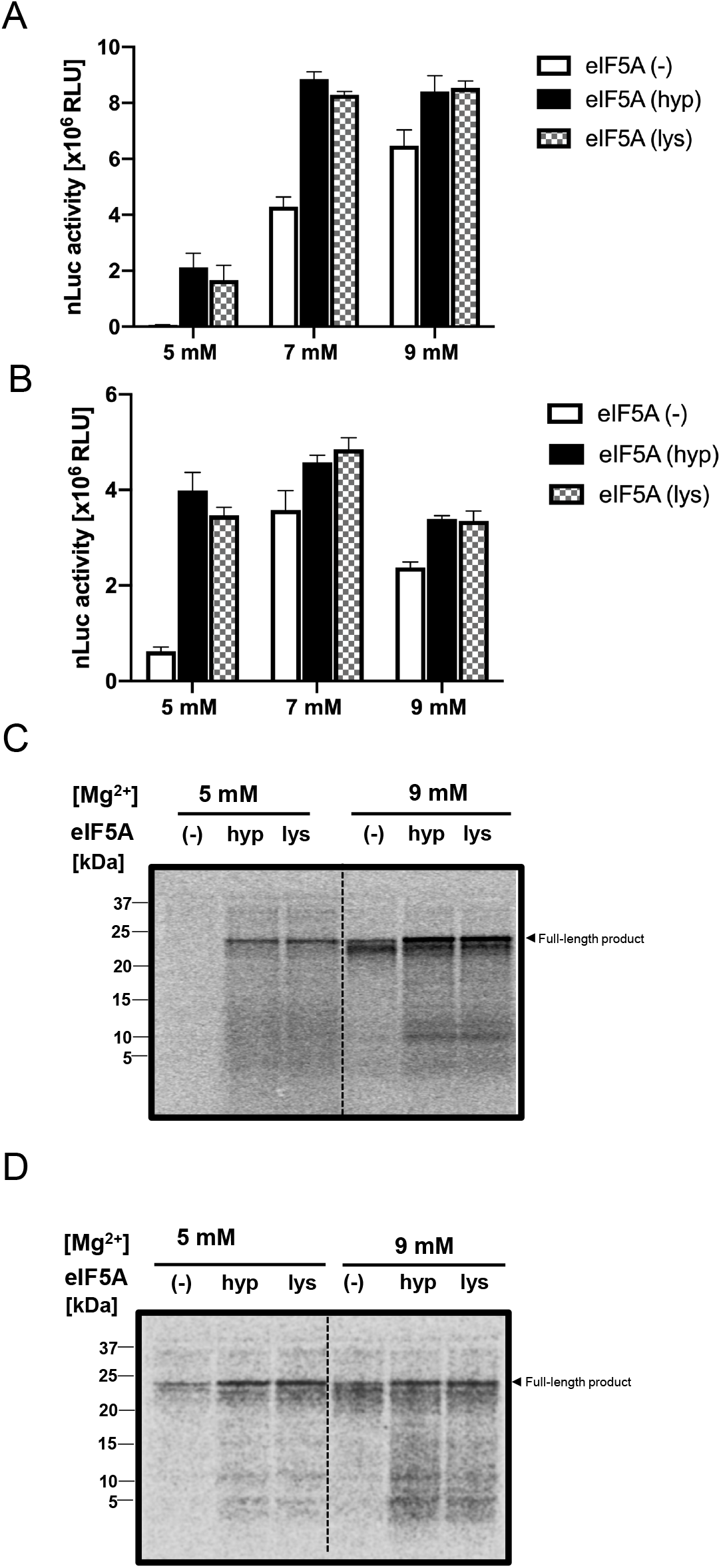
The effect of different concentrations of Mg2+ on the synthesis of full-length NanoLuc in the absence or presence of 2.5 μM eIF5A (hyp) and eIF5A (lys). The NanoLuc activity was detected after 3 hr reaction. (A) The synthesis of NanoLuc with iVT aa-tRNAs (14.4 μM) and (B) native aa-tRNAs (7.2 μM) in the context of 5 mM, 7 mM and 9 mM Mg2+. n=3. (C) The Tricine SDS-PAGE electrophoresis of full-length product after 3hr reaction, with [35S] iVT aa-tRNAs and (D) [35S] native aa-tRNAs at the 5 mM Mg2+ and 9 mM Mg2+.

At the upper end of the Mg2+ range, however, product profiles broadened, and additional off-target bands appeared, indicating that accelerated elongation is accompanied by reduced fidelity (Fig. 4; Supp. Fig. 2). Thus, Mg2+ can independently promote long-ORF completion in this system, but high concentrations carry a measurable accuracy cost.

### eIF5A resolves early blocks and enriches RNase-resistant products, whereas high Mg2+ gives a cruder rescue

As a baseline control, reactions without mRNA produced no RNase-resistant peptide either with or without eIF5A; in the −RNase lanes a single signal corresponding to [35S] Met– tRNA was observed and vanished upon RNase treatment, confirming its assignment (Fig. 5A). With the full-length NanoLuc mRNA at standard Mg2+ and +RNase, reactions lacking eIF5A showed no RNase-resistant peptide, whereas adding eIF5A yielded discrete RNase-resistant short peptides, indicating that eIF5A enables progression beyond initiation into productive elongation (Fig. 5B). To test whether stalling arises from the N-terminal FLAG poly-proline motif, we removed this tag—ΔFLAG removes the N-terminal DYKDDDDK (poly-Asp-rich) tag—and repeated the assay. Under the same baseline conditions without eIF5A, ΔFLAG mRNA still failed to produce RNase-resistant peptides; in the −RNase lanes only tRNA-linked signal was detected, consistent with very early (initiation/early-elongation) arrest rather than a discrete poly-Asp stall (Fig. 5C). Finally, raising Mg2+ to 9 mM in the absence of eIF5A increased the RNase-resistant full-length band and revealed additional RNase-resistant products, demonstrating an eIF5A-independent route to promote elongation (Fig. 5D). However, the product pattern broadens at high Mg2+, consistent with a loss of translational fidelity compared to the cleaner eIF5A-driven profile (Fig. 5D). Where measured, endpoint NanoLuc activities tracked these assignments, increasing with +eIF5A and with 9 mM Mg2+ relative to baseline, while the eIF5A condition yielded a cleaner product distribution than high Mg2+.

**Figure 5.**
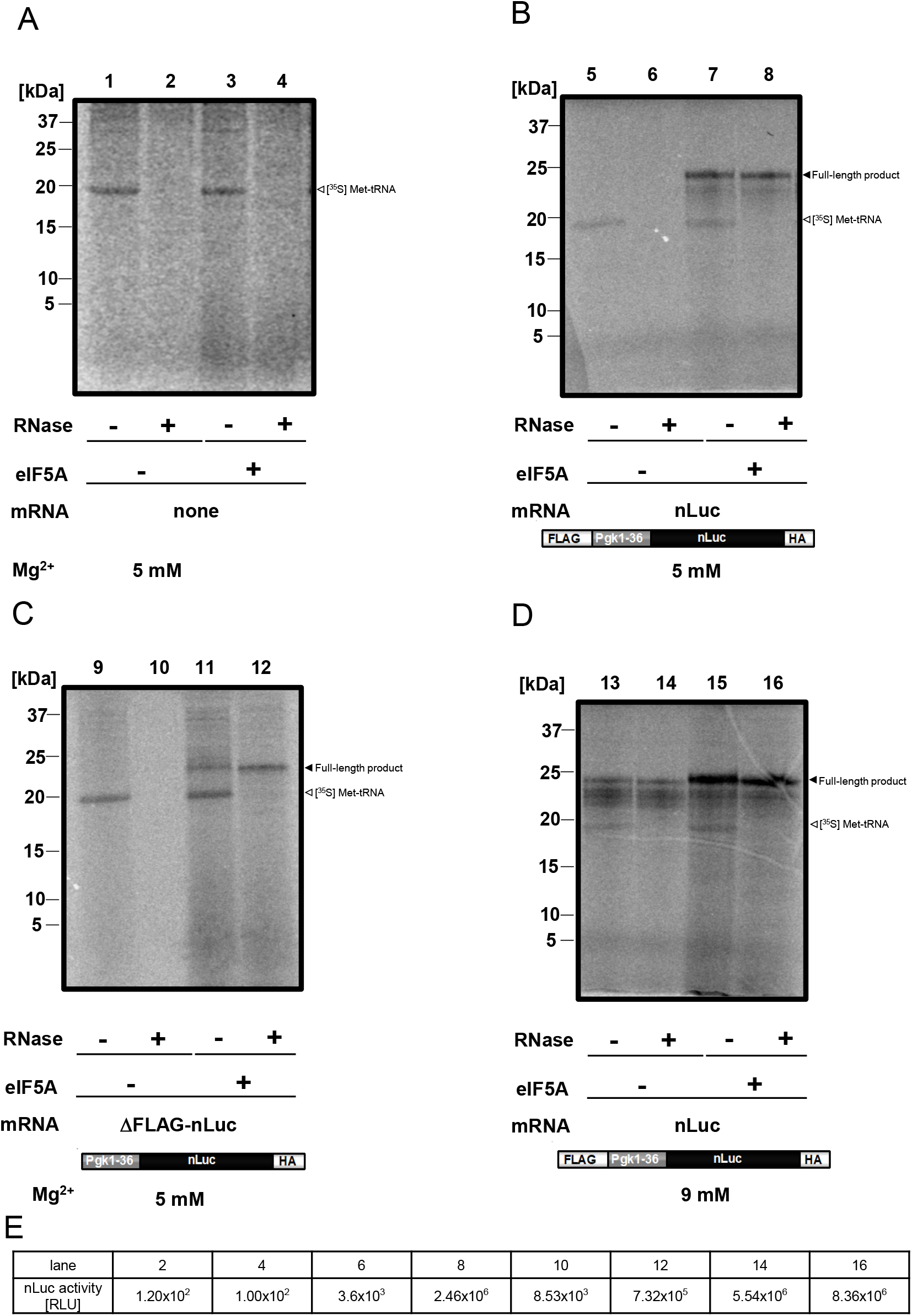
The Tricine SDS-PAGE electrophoresis of NanoLuc synthesis, using iVT aa-tRNAs (14.4 μM) in the absence or presence of 2.5 μM eIF5A (hyp). After 3hr reaction, the systems were treated with RNase A or not. (A) The electrophoresis without mRNA. (B) The electrophoresis of full-length NanoLuc in the context of NanoLuc mRNA at 5 mM Mg2+. (C) The electrophoresis of full-length NanoLuc in the context of NanoLuc mRNA without Flag sequence at 5 mM Mg2+. (D) The electrophoresis of full-length NanoLuc in the context of NanoLuc mRNA at 9 mM Mg2+. (E) The NanoLuc activities of respective lanes in (A-D), detected after 3hr reaction.

### Core eIF5A residues R27 and R87 underpin rescue of long-ORF translation and relief of poly-Pro stalling

Because hypusination was not required for rescue in the IVT-tRNA system, we tested whether contacts from the eIF5A body to the P-site tRNA contribute to activity. Guided by the eIF5A—P-site tRNA interface (model from PDB 5GAK), we focused on two basic residues-R27 and R87-that lie at the protein – tRNA contact surface (Fig. 6A). We constructed R27E and R87E single mutants in the lysine form of eIF5A [eIF5A(lys)], as well as the R27E/R87E double mutant, and assayed full-length NanoLuc synthesis at 6 mM Mg2+ with IVT aa-tRNAs (14.4 μM) and 2.5 μM eIF5A. Relative to wild-type eIF5A(lys) (set to 100%), each single substitution caused a modest reduction, whereas the double mutant markedly abrogated the eIF5A-dependent promotion of long-ORF translation (Fig. 6B). Time-course measurements at the same conditions corroborated this ranking: eIF5A(lys) (and eIF5A(hyp)) drove earlier onset and higher accumulation of full-length product, while R27E/R87E showed delayed and lower plateaus (Supp. Fig. 3).

**Figure 6.**
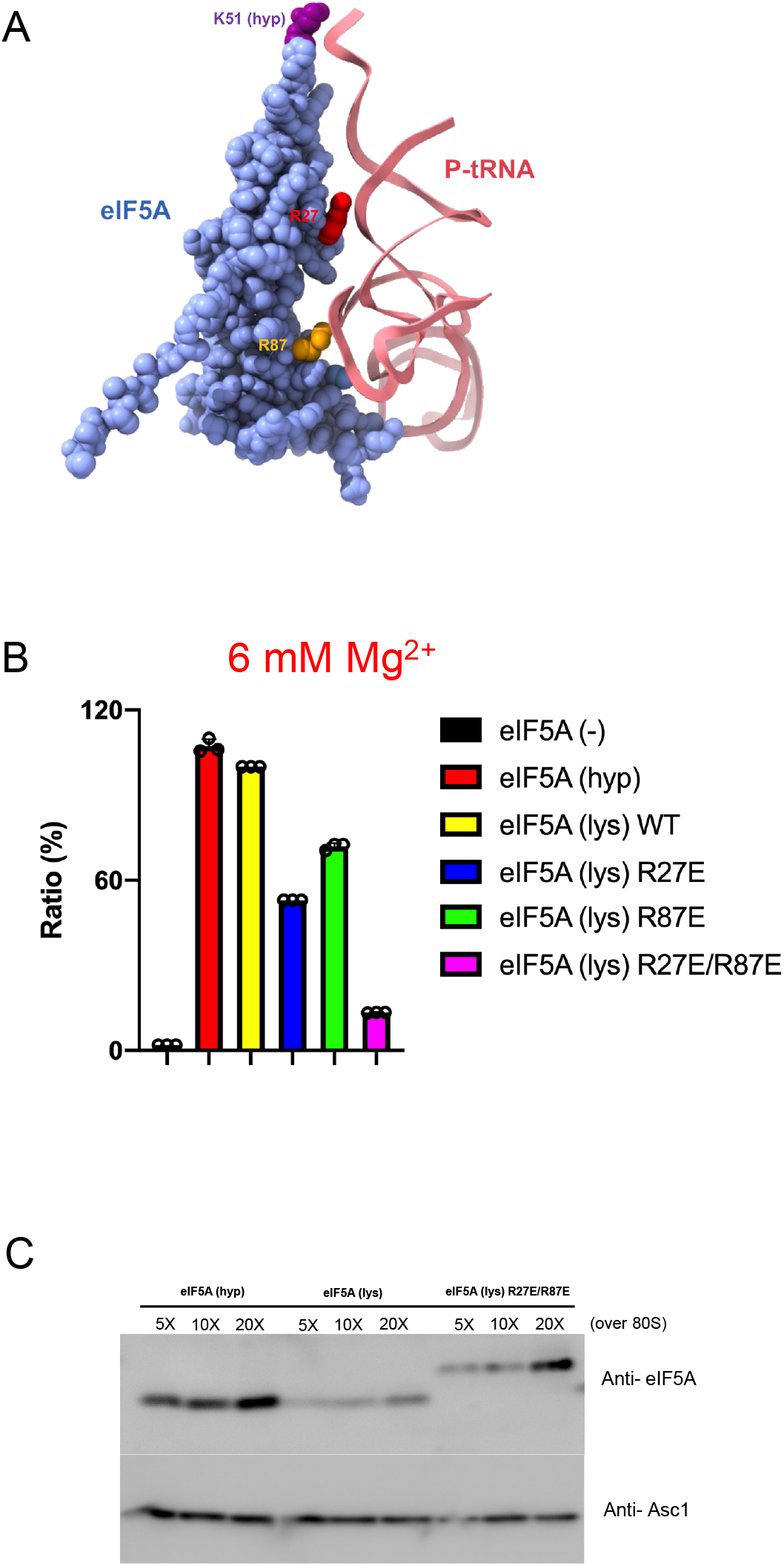
The effect of eIF5A (lys) mutants on the synthesis of full-length NanoLuc with iVT aa-tRNAs at 6 mM Mg2+. (A) The schematic structure of eIF5A interacting with P-site tRNA (from PBK: 5gak). The blue part is the body of eIF5A. The pink part is P-site tRNA. The violet part is hypusinated lysine of eIF5A. The red part is the 27th amino acid (lysine) on the body of eIF5A and the yellow part is the 87th amino acid (lysine). (B) The synthesis ratio of NanoLuc with iVT aa-tRNAs (14.4 μM) in the presence of different types of eIF5A (2.5 μM), in comparison with wild eIF5A (lys) (100%). n=3. (C) The Western Blot of eIF5A and mutants binding with the yeast ribosome. The Asc1 was considered as the ribosome control. The concentration folds of eIF5A were compared to the constant concentration of ribosome (0.5 μM) in the reaction system.

To distinguish loss of catalysis from loss of recruitment, we assessed 80S binding by immunoblot while holding ribosomes at 0.5 μM. eIF5A(hyp) displayed strong binding even at the lower protein inputs, and eIF5A(lys) and R27E/R87E reached similar binding levels at higher inputs (Fig. 6C). Thus, the functional deficit of R27E/R87E is not explained by impaired ribosome association but is consistent with weakened P-site tRNA stabilization/peptidyl-transfer support provided by the eIF5A body.

We next asked whether these body residues are also required to relieve poly-proline (poly-Pro) stalls, a canonical eIF5A phenotype. In reactions with native tRNAs, eIF5A(lys) alleviated the elongation block imposed by an internal poly-Pro motif, whereas the R27E/R87E mutant failed to relieve this stall (Fig. 7B) and did not enhance translation of a control mRNA lacking the motif (Fig. 7A). Taken together, Figures 6–7 (with Supp. Fig. 3) show that R27 and R87 in the eIF5A core are essential for eIF5A-mediated promotion of long-ORF synthesis and for relief of poly-Pro–induced pausing, while ribosome binding per se is largely preserved. These findings, together with the robust charging of “naked” IVT tRNAs, suggest that tRNA modifications and eIF5A body contacts act in concert at the P site to stabilize the tRNA and support efficient peptidyl transfer in the eukaryotic system.

**Figure 7.**
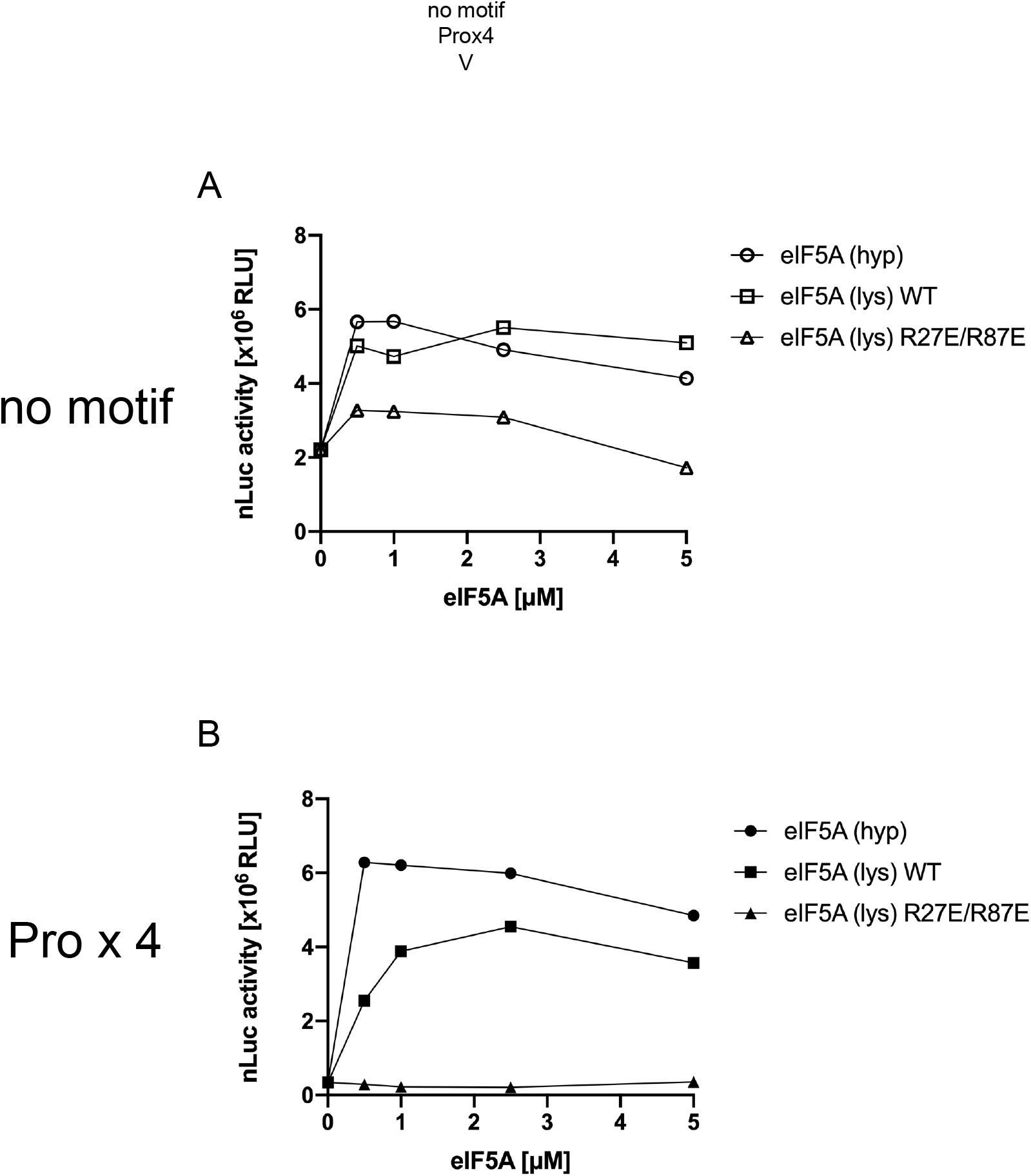
The effect of eIF5A (lys) dual-site mutant on the translation of full-length NanoLuc mRNA and [ProX4] inserted mRNA with native aa-tRNAs (7.2 μM) at 5mM Mg2+. The mRNA sequences were designed according to our previous article. (A) The translation of NanoLuc mRNA and (B) [ProX4] inserted mRNA with native aa-tRNAs, in the presence of 2.5 μM eIF5A (hyp), eIF5A (lys) and eIF5A (lys) R27E/R87E mutant.

### Mg2+ accelerates translation but increases stop-codon readthrough while eIF5A preserves fidelity

Gel analyses at 9 mM Mg2+ revealed additional products below the full-length band, suggesting mistranslation. To quantify accuracy, we used readthrough reporters bearing a single UAA/UGA/UAG upstream of NanoLuc and defined readthrough as the ratio of luminescence from the stop-inserted mRNA to that from the matched sense control. We first fixed the assay window at 1.5 h, which lay in the linear regime with respect to mRNA dose and time across conditions (Supp. Fig. 5).

At 5 mM Mg2+ with eRF1 present, readthrough was ∼1–3% in both native-tRNA and IVT-tRNA reactions (Fig. 8A,B,E; Supp. Fig. 4A,C,I). Raising Mg2+ to 9 mM increased readthrough to ∼10–15% and this elevation was independent of eIF5A (Fig. 8C,D,F,G; Supp. Fig. 4E,G,K,M). In the absence of eRF1, readthrough occurred even with native tRNAs and became markedly stronger at 9 mM Mg2+ (Supp. Fig. 4B, D, F, H, J, L, N), underscoring the requirement for termination factors.

**Figure 8.**
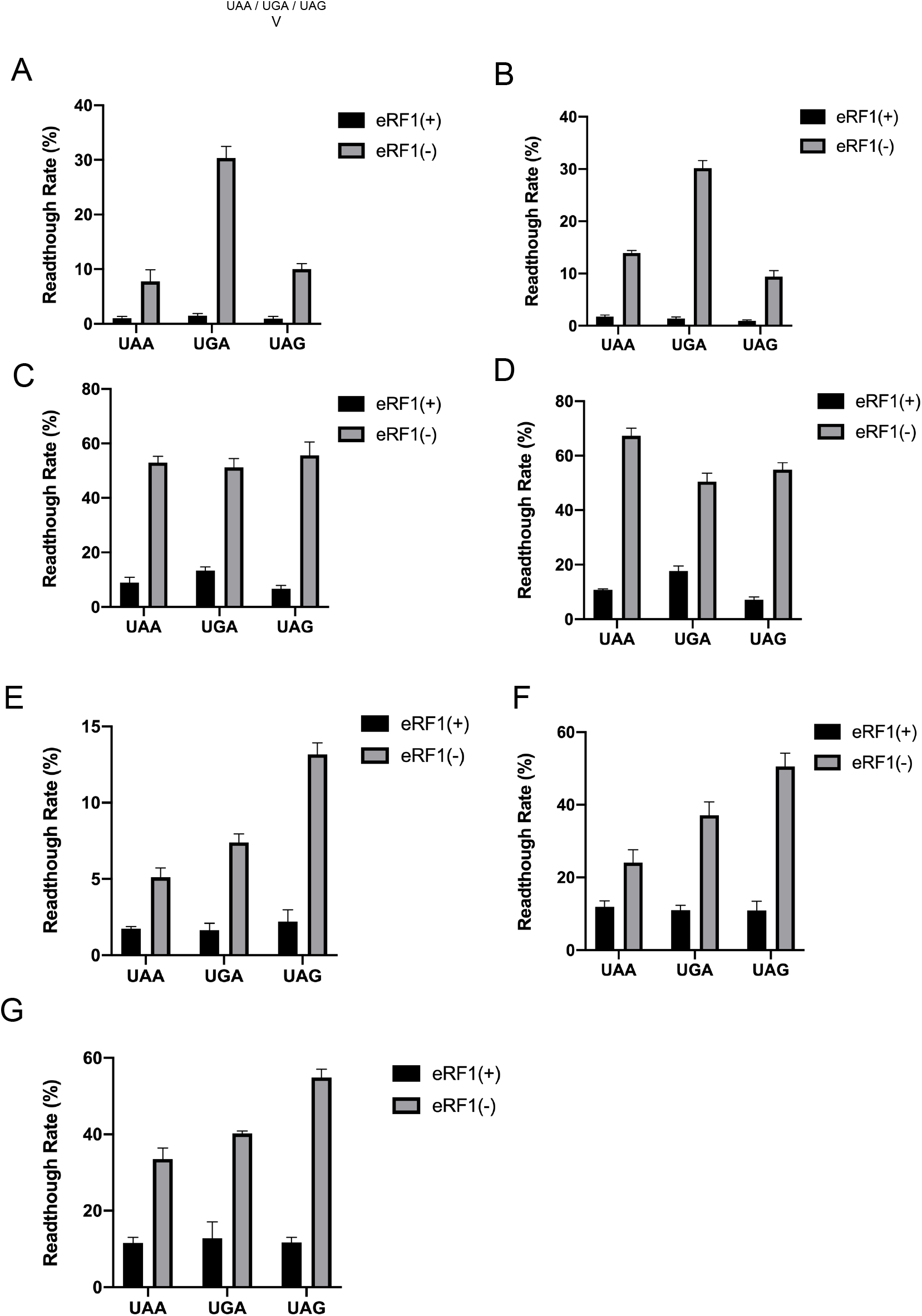
Increasing Mg2+ levels in the yeast PURE system can improve the readthrough efficiency. The rates were calculated by the ratio of stop codons inserted mRNA with the normal full-length NanoLuc mRNA (no motif), in the absence or presence of eRF1(0.5 μ M). The NanoLuc activities were collected after 1.5hr reaction. n=3. (A) The readthrough rates with native aa-tRNAs (7.2 μM) in the absence of eIF5A (hyp) at the 5 mM Mg2+. (B) The readthrough rates with native aa-tRNAs in the presence of 2.5 μM eIF5A (hyp) at the 5 mM Mg2+. (C) The readthrough rates with native aa-tRNAs in the absence of eIF5A (hyp) at the 9 mM Mg2+. (D) The readthrough rates with native aa-tRNAs in the presence of 2.5 μM eIF5A (hyp) at the 9 mM Mg2+. (E) The readthrough rates with iVT aa-tRNAs (14.4 μM) in the presence of 2.5 μM eIF5A (hyp) at the 5 mM Mg2+. (F) The readthrough rates with iVT aa-tRNAs (14.4 μM) in the absence of eIF5A (hyp) at the 9 mM Mg2+. (G) The readthrough rates with iVT aa-tRNAs (14.4 μM) in the presence of 2.5 μM eIF5A (hyp) at the 9 mM Mg2+.

Stop-codon preferences differed between pools. With native tRNAs at 5 mM Mg2+, UGA > UAA ≈ UAG (Fig. 8A,B), whereas at 9 mM this bias was largely lost (Fig. 8C,D). With IVT tRNAs, the ordering UAG > UGA > UAA held at both 5 mM and 9 mM Mg2+ (Fig. 8E–G). Thus, while eIF5A supports efficient long-ORF synthesis without increasing readthrough, Mg2+ provides a broad, factor-independent boost to elongation at the cost of fidelity. The higher suppressor activity and altered stop-codon preference seen with IVT (minimally modified) tRNAs further indicate that tRNA modifications contribute to eukaryotic translational fidelity beyond their roles in peptidyl transfer.

## Discussion

We established a fully defined S. cerevisiae system that relies on a minimal pool of aminoacylated IVT tRNAs. Long ORFs translate efficiently when either eIF5A is supplied or Mg2+ is elevated, whereas short ORFs translate similarly with IVT or native tRNAs. Importantly, eIF5A enhances yield without increasing readthrough, while high Mg2+ speeds elongation at the expense of fidelity (Figs. 2–5, 8; Supp. Figs. 1, 4–5).

Classic and recent work assigns eIF5A to difficult elongation steps and specific sequence contexts: discovery/definition(Kemper, Berry & Merrick, 1976; Safer, 1989), abundance (Hukelmann et al., 2016), structure (Melnikov et al., 2016), hypusination pathway and biology (Park et al., 2010), initiation/early peptide-bond assays (Gregio et al., 2009), relief of poly-proline stalls(Gutierrez et al., 2013), ribosome binding and positioning near the E/P sites (Doerfel et al., 2015). Here, hypusination is not required for rescue in vitro: eIF5A(hyp) and eIF5A(lys) perform similarly at micromolar levels (Figs. 2–3; Supp. Fig. 1). Structure-guided mutagenesis shows that R27/R87 in the eIF5A body are critical: R27E/R87E markedly reduces full-length NanoLuc while 80S binding remains comparable to wild type (Fig. 6; Supp. Fig. 3), placing the defect at P-site tRNA stabilization/peptidyl-transfer support rather than recruitment. Consistently, R27E/R87E fails to relieve internal poly-Pro stalling in native-tRNA reactions (Fig. 7), in line with genetic/biochemical evidence that eIF5A promotes difficult peptide bonds and can influence start-codon selection and termination (Manjunath et al., 2019).

As a complementary lever, raising Mg2+ (5→9 mM) restores long-ORF completion without eIF5A (Figs. 4–5) but increases stop-codon readthrough from ∼1–3% to ∼10–15% at 1.5 h, regardless of eIF5A, and the effect becomes stronger without eRF1 (Fig. 8; Supp. Fig. 4). These data fit models in which Mg2+—pervasive across ribosomal architecture, factor cycles, and tRNA tertiary structure (Xu, MacKerell & Nilsson, 2016; Yamagami et al., 2018; Yamagami, Sieg & Bevilacqua, 2021)—stabilizes near/non-cognate interactions and broadens decoding acceptance, thereby speeding elongation while reducing accuracy (Shenvi et al., 2005; Yamamoto et al., 2010).

Notably, stop-codon preferences differ between pools: with native tRNAs at 5 mM Mg2+ we observe UGA > UAA ≈ UAG, whereas IVT tRNAs show UAG > UGA > UAA at both Mg2+ levels (Fig. 8). The higher suppressor activity and altered ordering with IVT tRNAs point to missing nucleotide modifications as contributors to eukaryotic decoding specificity, not only to peptidyl-transfer efficiency. This is consistent with prior work on tRNA tertiary stabilization and modification–Mg2+ synergy(Xu, MacKerell & Nilsson, 2016; Schauss et al., 2021) and with functional studies on t6A37 and m1G37 in decoding fidelity/frameshift control (Deutsch et al., 2012; El Yacoubi, Bailly & De Crécy-Lagard, 2012; Hou, Masuda & Gamper, 2019).

Together, these results separate factor-specific and ionic routes to completing long ORFs in a defined eukaryotic setting: eIF5A—via body residues such as R27/R87—promotes elongation while preserving accuracy, whereas elevated Mg2+ provides a broad accelerant at a fidelity cost. The minimal 21-member IVT-tRNA pool makes the system highly programmable; in future, systematic add-back of defined tRNA features and codon-context panels, coupled to time-resolved termination assays, toeprinting/ribosome-profiling, and MS verification of products, should deconvolve contributions to P-site stabilization vs. decoding under controlled ionic conditions. Although our measurements were made in vitro at fixed factor/Mg2+/tRNA concentrations, the agreement with established eIF5A functions and Mg2+ effects supports generality. Practically, the platform offers a rapid benchmark for genetic-code engineering (e.g., orthogonal pairs, release-factor variants) while providing quantitative readouts of the speed–fidelity trade-off.

## Methods And Materials

### Plasmids and reporter templates

Yeast tRNA genes (Supp. Table 1) were PCR-amplified with primers adding a T7 promoter and cloned into pCR4-TOPO (Invitrogen, Thermo Fisher Scientific; Waltham, MA, USA) at EcoRI sites; ligation products were transformed into XL10-Gold (Agilent Technologies; Santa Clara, CA, USA) and sequence-verified. eIF5A variants were generated by PCR-based site-directed mutagenesis on pET-29b (Novagen, Merck KGaA; Darmstadt, Germany) and confirmed by Sanger sequencing. Reporter DNAs included NanoLuc, ΔFLAG (N-terminal DYKDDDDK removed), single premature-termination reporters (UAA/UGA/UAG inserted upstream of NanoLuc), and internal poly-proline insertions. (Supp. Table 2)

### IVT tRNA production and aminoacylation

tRNA genes were transcribed in vitro with T7 RNA polymerase. Transcripts requiring 5′-end maturation (tRNA^i^Met_CAU_, tRNAAsp_GUC_, tRNAGlu_CUC_, tRNATyr_GUA_) were trimmed using E. coli RNase P. All IVT tRNAs were refolded, purified by HiTrap Q HP (Cytiva; Marlborough, MA, USA) anion-exchange, and assessed by denaturing PAGE. Each tRNA was aminoacylated individually using either a yeast S100 preparation or recombinant aaRS (e.g., *E. coli* MetRS, LysRS, GlnRS; yeast ProRS). Charging efficiencies were quantified by incorporation of radiolabeled amino acids (PerkinElmer; Waltham, MA, USA) followed by liquid scintillation counting (selected species corroborated by acid-urea PAGE shifts). For translation, IVT aa-tRNAs were pooled to 14.4 μM (native aa-tRNA pools, where used, 7.2 μM).

*Note on S100*: YPH499 cells were grown in YPD and mechanically disrupted on ice; lysates were clarified at 20,000 × g and 100,000 × g to obtain S100, dialyzed and polished on Q-Sepharose (Cytiva; Marlborough, MA, USA), then stored at −80 °C.

### mRNA synthesis and purification

PCR-amplified templates were transcribed with T7 RNA polymerase. mRNAs were purified by phenol–chloroform (FUJIFILM Wako Pure Chemical Corporation; Osaka, Japan) extraction, ethanol precipitation, and desalting on MicroSpin columns (Cytiva; Marlborough, MA, USA), quantified by A260, and stored at −80 °C.

### Defined yeast translation reactions

Reactions contained purified yeast 80S ribosomes, elongation factors (eEF1A, eEF2, eEF3), release factors (eRF1, eRF3), aminoacylated IVT (or native) tRNA pools, and synthetic mRNA. Unless stated otherwise, incubations were at 30 °C for 3 h. Mg2+ was titrated as indicated (typically 5 mM vs 9 mM). eIF5A supplements—hypusinated eIF5A(hyp) or lysine form eIF5A(lys)—were added where noted (typically 2.5 μM).

### Fidelity (stop-codon readthrough) assay

Single-stop reporters (UAA/UGA/UAG) were translated in parallel with the matched sense control. To standardize rate effects, readthrough was quantified at 1.5 h, which lies in the linear regime with respect to mRNA dose and time; eRF1 was present at 0.5 μM or omitted as specified.

Readthrough (%) = *(luminescence of stop reporter at 1*.*5 h) / (luminescence of sense control at 1*.*5 h) × 100*.

### Protein expression, purification, and ribosome binding

Wild-type and mutant eIF5A were expressed in Rosetta (DE3) induced by IPTG, purified by HiTrap Q HP (Cytiva; Marlborough, MA, USA), dialyzed, aliquoted, and stored at −80 °C; both eIF5A(hyp) and eIF5A(lys) were used as indicated. For 80S binding assays, reactions contained a constant ribosome concentration (0.5 μM) with titrated eIF5A; complexes were recovered by sucrose-cushion ultracentrifugation and analyzed by SDS-PAGE/Western blot with anti-eIF5A; Asc1 served as a ribosomal control.

### Product detection and data analysis

Translation products were resolved by Tricine–SDS-PAGE. For radiolabeling experiments, [35S]-methionine (PerkinElmer; Waltham, MA, USA) was included and products visualized by autoradiography. To distinguish peptidyl-tRNA from polypeptide, matched samples were processed ±RNase A (Sigma-Aldrich, Merck KGaA; Darmstadt, Germany): RNase-resistant bands correspond to peptide products, whereas RNase-sensitive signals reflect peptidyl-tRNA or [35S] Met–tRNA species. Enzymatic activity was measured with Nano-Glo on GloMax luminometers (Promega, Madison, WI, USA). Unless otherwise specified, data are mean ± SD from ≥3 independent experiments; *n* and statistical tests are reported in figure legends. Gel densitometry used local background subtraction and normalization to match controls.

## Figure Legend

Supplementary Figure 1. The time course of the synthesis reaction under 5 mM Mg2+ in the presence of different concentrations of eIF5A (hyp) and eIF5A (lys), detected at the 1st hr, 2nd hr and 3rd hr.

(A)and (B) The synthesis of HiBiT with iVT aa-tRNA (14.4 μM) and native aa-tRNA (7.2 μ M).

(C) and (D) The synthesis of full-length NanoLuc with iVT aa-tRNA and native aa-tRNA.

Supplementary Figure 2. The time course of the synthesis of full-length NanoLuc at different concentrations of Mg2+ in the absence or presence of 2.5 μM eIF5A (hyp) and eIF5A (lys).

(A)The synthesis of NanoLuc at 5 mM, 7 mM and 9 mM Mg2+ with iVT aa-tRNAs, in the absence of eIF5A or the presence of (B) 2.5 μM eIF5A (hyp) and (C) eIF5A (lys).

(D) The synthesis of NanoLuc at 5 mM, 7 mM and 9 mM Mg2+ with native aa-tRNAs, in the absence of eIF5A or the presence of (E) 2.5 μM eIF5A (hyp) and (F) eIF5A (lys).

Supplementary Figure 3. The time course of synthesis of full-length NanoLuc with iVT aa-tRNAs (14.4 μM) in the absence or precence of different types of eIF5A (2.5 μM) at 6 mM Mg2+.

Supplementary Figure 4. The time courses of readthrough assays in Figure 8. The NanoLuc activities were collected after 1.5hr and 3hr reaction.

(A)The NanoLuc activities in the context of native aa-tRNAs (7.2 μM) in the absence of eIF5A (hyp) at the 5 mM Mg2+, with eRF1 or (B) without eRF1.

(C) The NanoLuc activities in the context of native aa-tRNAs (7.2 μM) in presence of 2.5 μM eIF5A (hyp) at the 5 mM Mg2+, with eRF1 or (D) without eRF1.

(E) The NanoLuc activities in the context of native aa-tRNAs (7.2 μM) in the absence of

2.5 μM eIF5A (hyp) at the 9 mM Mg2+, with eRF1 or (F) without eRF1.

(G) The NanoLuc activities in the context of native aa-tRNAs (7.2 μM) in presence of 2.5 μM eIF5A (hyp) at the 9 mM Mg2+, with eRF1 or (H) without eRF1.

Supplementary 5. The correlation analysis of NanoLuc activities with input mRNA amount and reaction time. The synthesis of full-length NanoLuc was performed with different concentrations of input mRNA (0.2 μM, 0.5 μM and 1 μM), in the presence of eRF1. The NanoLuc activities were detected after 1.5 hr and 3 hr reaction.

(A)The synthesis of NanoLuc with native aa-tRNAs (7.2 μM), in the absence and (B) in the presence of 2.5 μM eIF5A (hyp) at 5 mM Mg2+.

(C) The synthesis of NanoLuc with native aa-tRNAs (7.2 μM), in the absence and(D) in the presence of 2.5 μM eIF5A (hyp) at 9 mM Mg^2+^.

(E) The synthesis of NanoLuc with iVT aa-tRNAs (14.4 μM), in the absence of eIF5A (hyp) at 5 mM Mg^2+^.

(F) The synthesis of NanoLuc with iVT aa-tRNAs (14.4 μM), in the absence and (G) in the presence of 2.5 μM eIF5A (hyp) at 9 mM Mg^2+^.

## Notes

https://doi.org/10.6084/m9.figshare.28904456

### Competing Interest Statement

The authors have declared no competing interest.

### Summary of Updates

This revision updates the manuscript for clarity, consistency, and closer alignment between data and claims. Title, abstract, keywords, and text were revised for precision. We removed statements about installing t6A37 or m1G37 on IVT tRNAs, since no modification add-back data are included in this study. Introduction was tightened and its final sentence rewritten to reflect the actual scope: a fully defined yeast translation platform using 21 IVT tRNAs, and quantitative analysis of how eIF5A and Mg2+ shape elongation and fidelity.

