## Supplementary File for "Synergy between eIF5A and Mg2+ enhances elongation in a defined yeast cell-free translation system with synthetic tRNAs"

### Slide 1
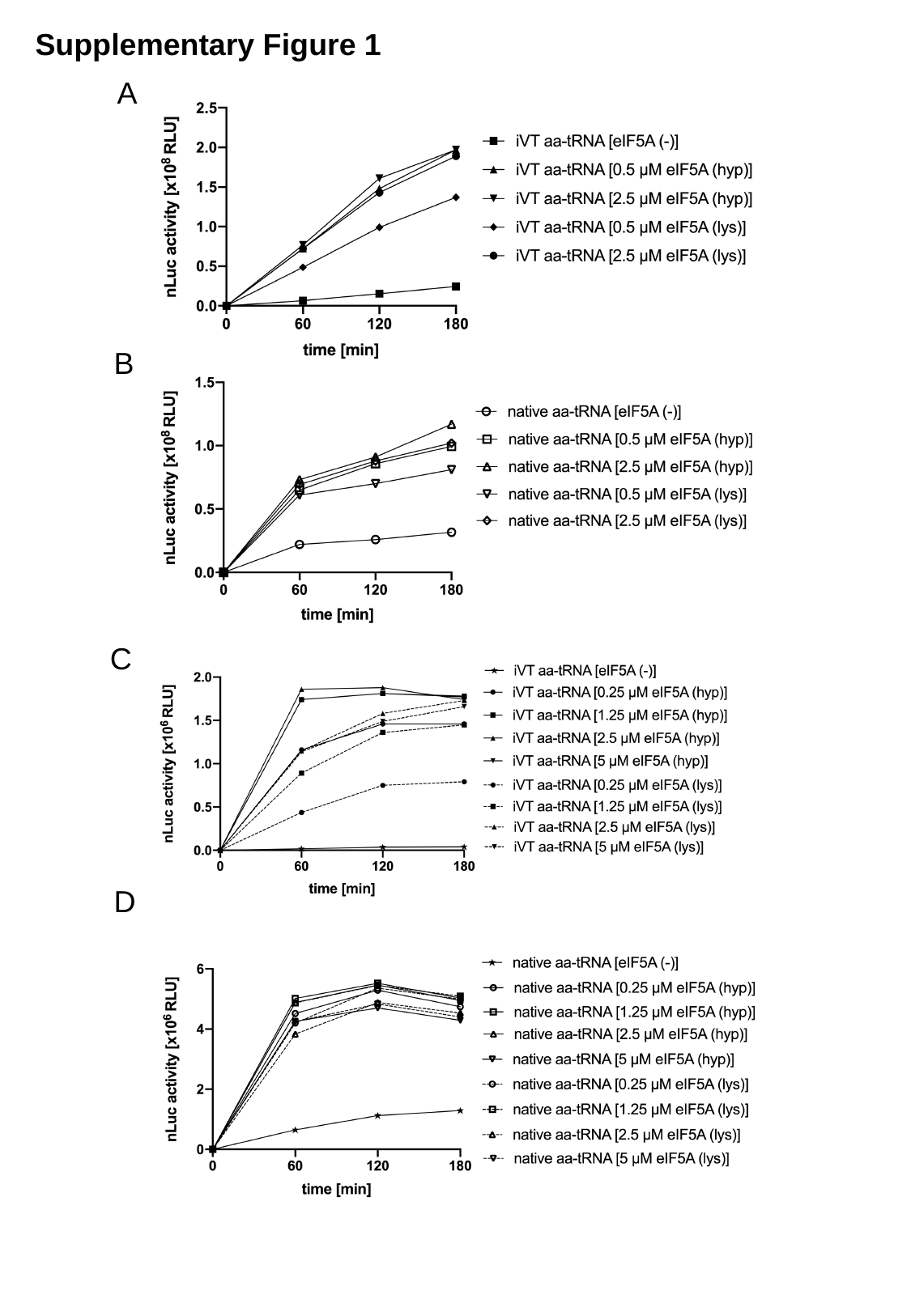

Supplementary Figure 1
A
B
C
D

### Slide 2
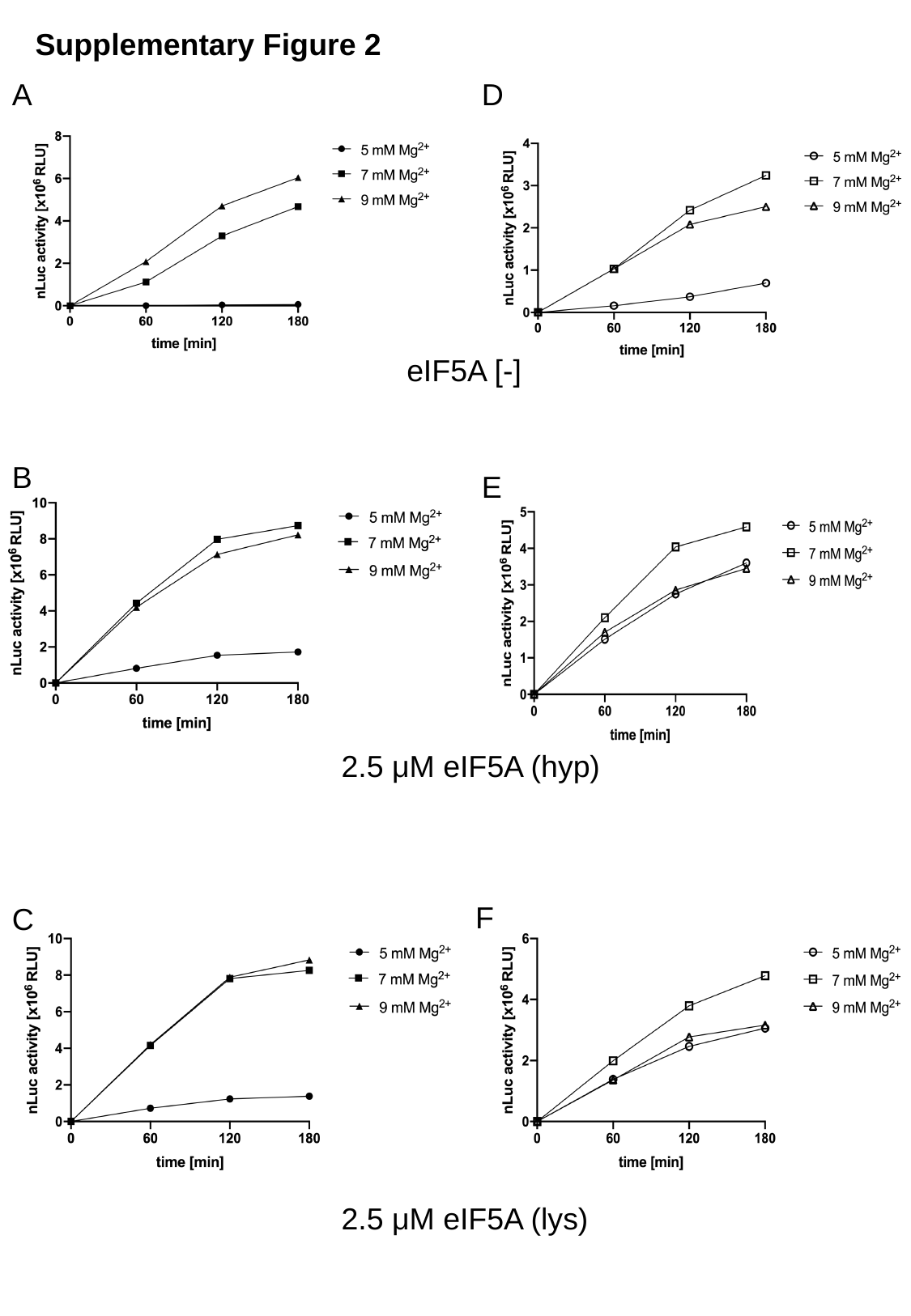

Supplementary Figure 2
A
D
eIF5A [-]
B
E
2.5 μM eIF5A (hyp)
F
C
2.5 μM eIF5A (lys)

### Slide 3
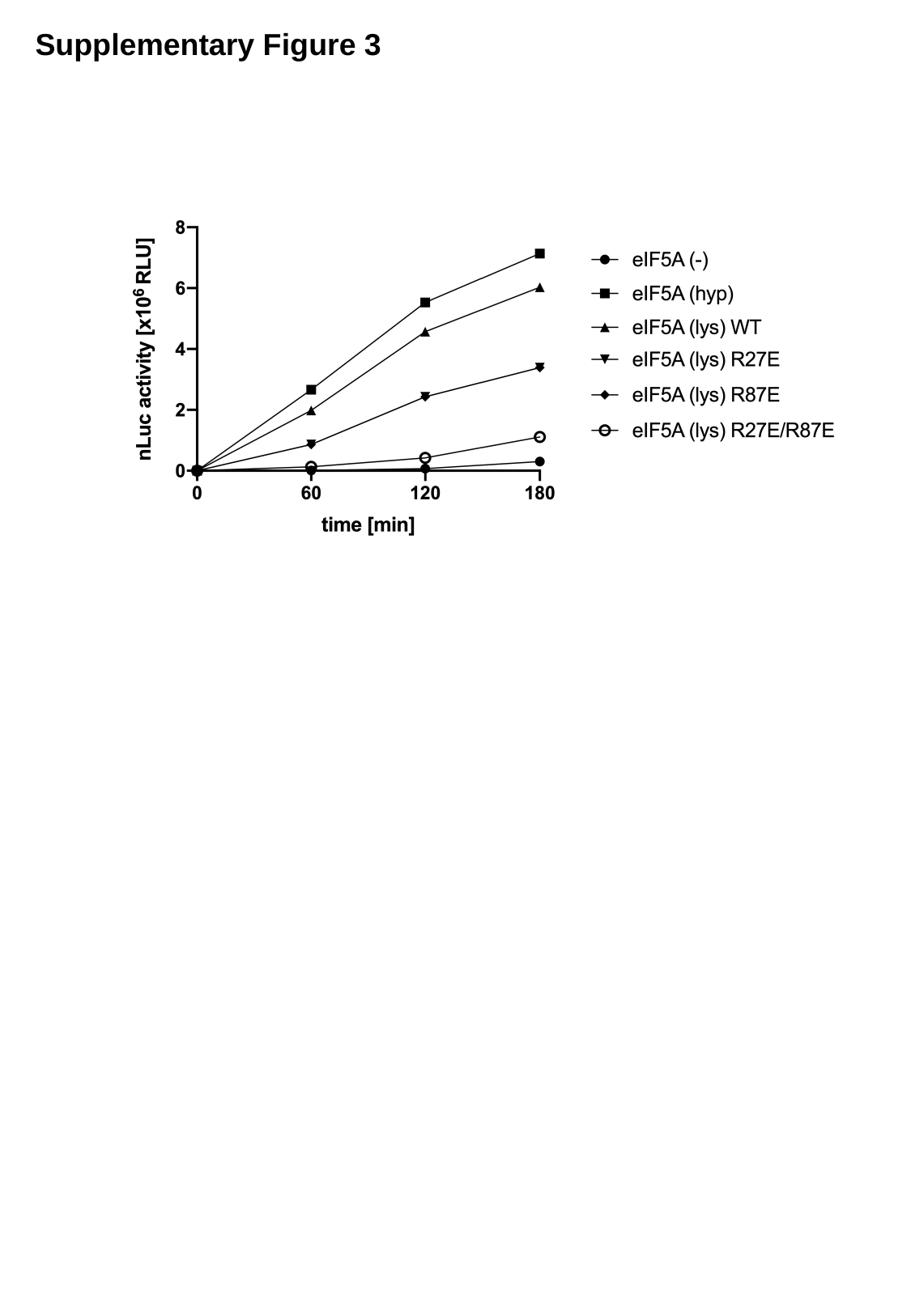

Supplementary Figure 3

### Slide 4
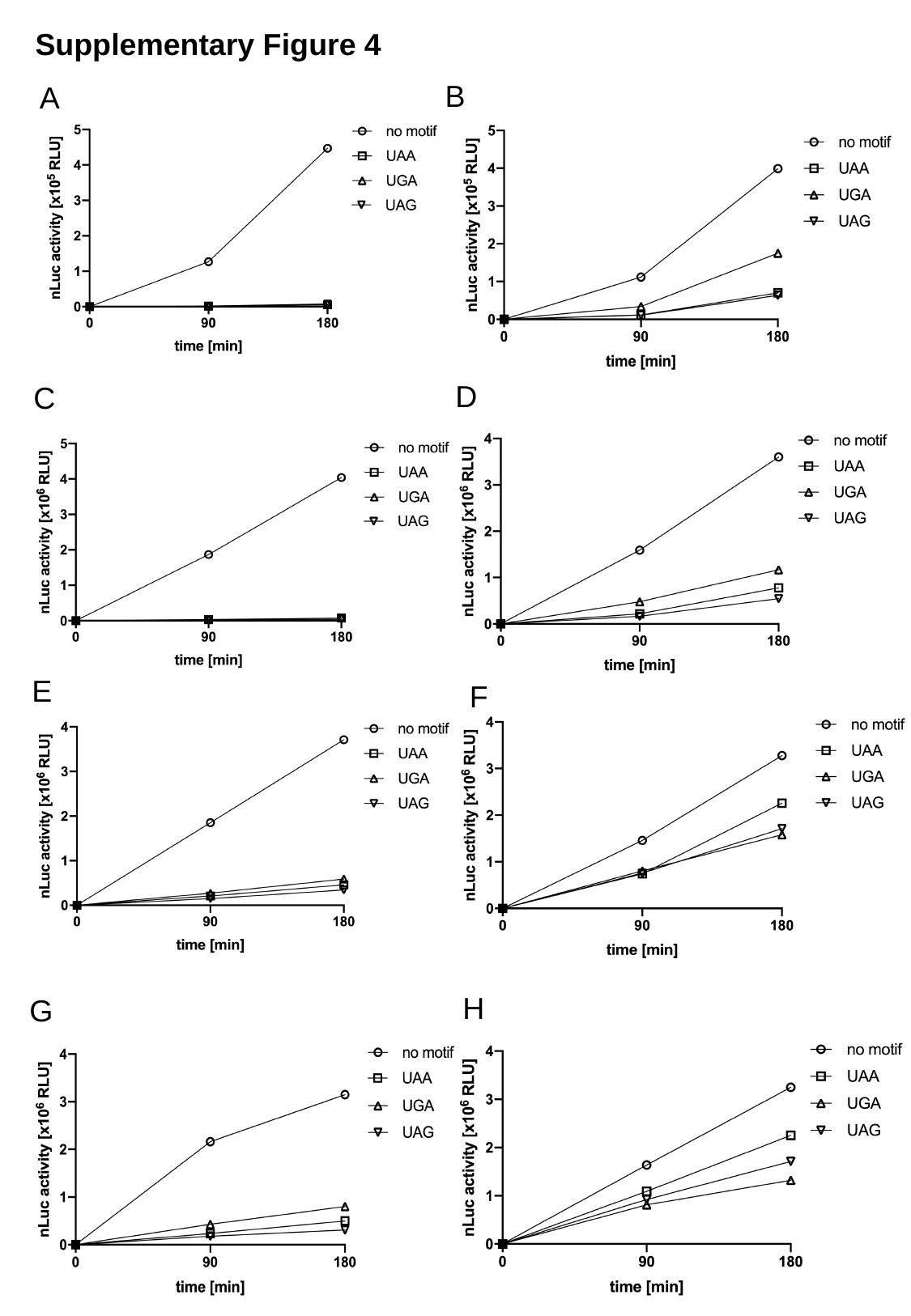

Supplementary Figure 4
B
A
D
C
E
F
H
G

### Slide 5
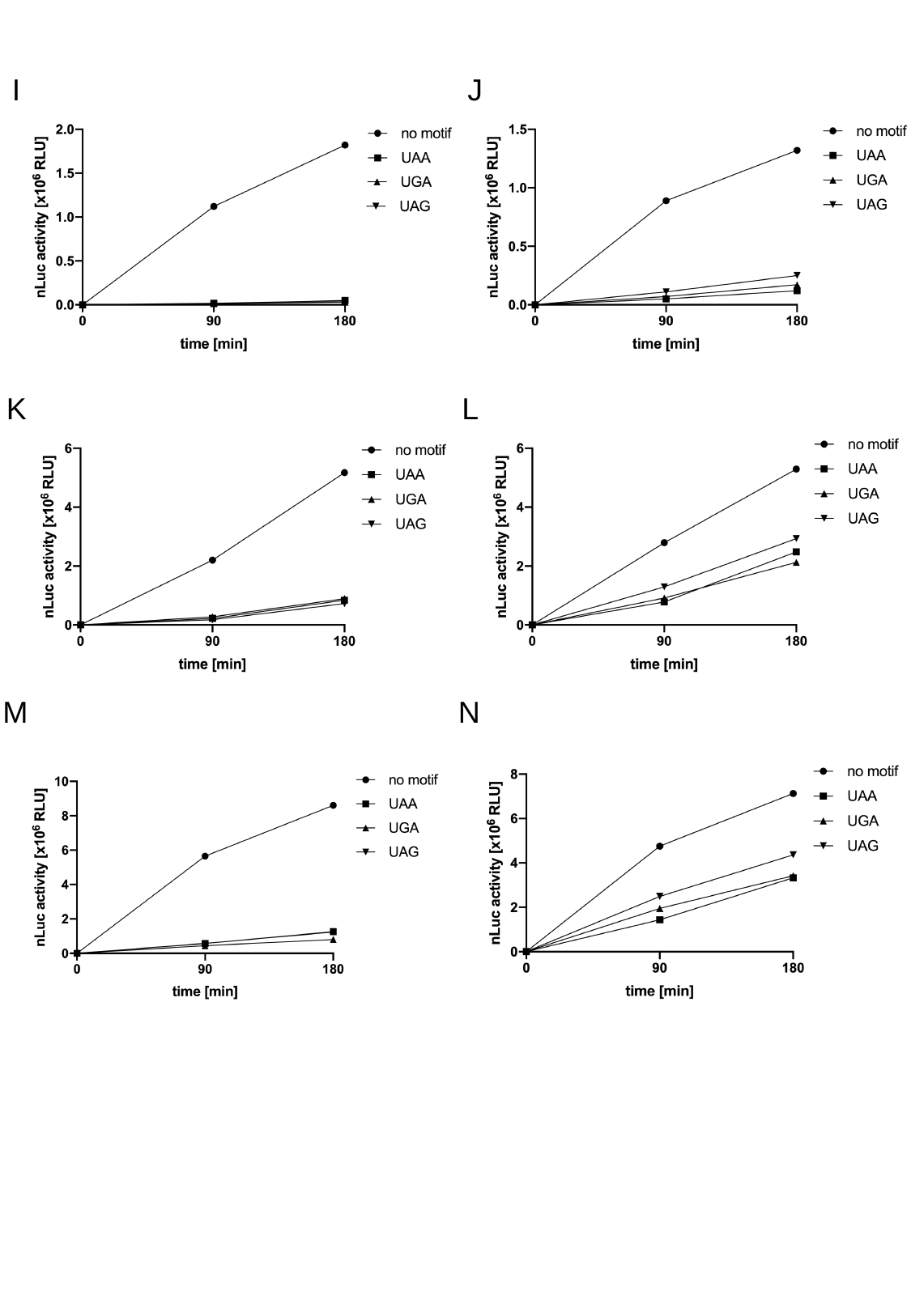

Supplementary Figure 4. The time courses of readthrough assays in Figure 8. The nanoluciferase activities were collected after 1.5hr and 3hr reaction. (A) The nanoluciferase activities in the context of native aa-tRNAs (7.2 μM) in the absence of eIF5A (hyp) at the 5 mM Mg^2+^, with eRF1 or (B) without eRF1. (C) The nanoluciferase activities in the context of native aa-tRNAs (7.2 μM) in presence of 2.5 μM eIF5A (hyp) at the 5 mM Mg^2+^, with eRF1 or (D) without eRF1. (E) The nanoluciferase activities in the context of native aa-tRNAs (7.2 μM) in the absence of 2.5 μM eIF5A (hyp) at the 9 mM Mg^2+^, with eRF1 or (F) without eRF1. (G) The nanoluciferase activities in the context of native aa-tRNAs (7.2 μM) in presence of 2.5 μM eIF5A (hyp) at the 9 mM Mg^2+^, with eRF1 or (H) without eRF1. (I) The nanoluciferase activities in the context of iVT aa-tRNAs (14.4 μM) in presence of 2.5 μM eIF5A (hyp) at the 5 mM Mg^2+^, with eRF1 or (J) without eRF1. (K) The nanoluciferase activities in the context of iVT aa-tRNAs (14.4 μM) in absence of eIF5A (hyp) at the 9 mM Mg^2+^, with eRF1 or (L) without eRF1. (M) The nanoluciferase activities in the context of iVT aa-tRNAs (14.4 μM) in presence of 2.5 μM eIF5A (hyp) at the 9 mM Mg^2+^, with eRF1 or (N) without eRF1.
I
J
K
L
M
N

### Slide 6
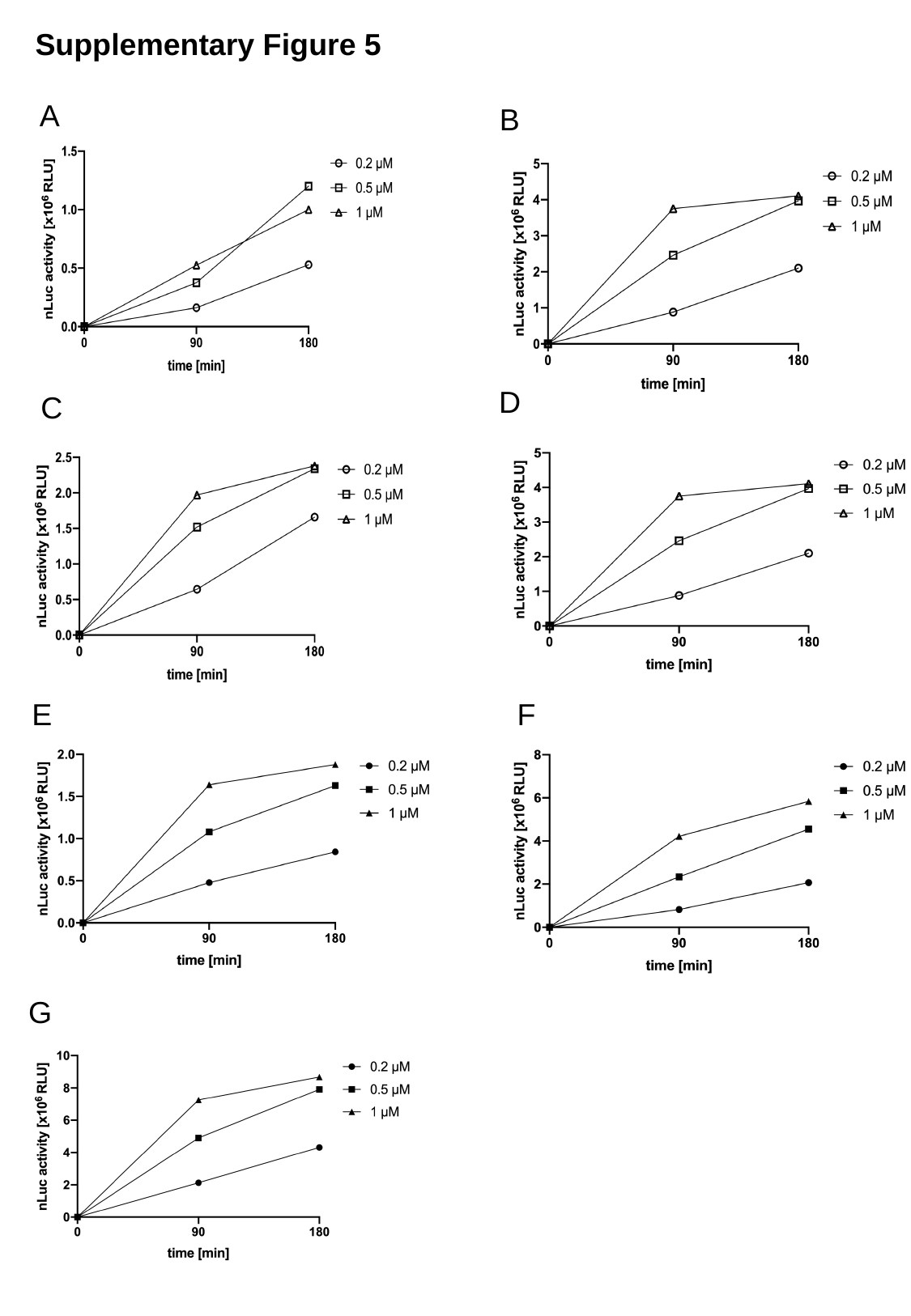

Supplementary Figure 5
A
B
D
C
E
F
G

### Slide 7
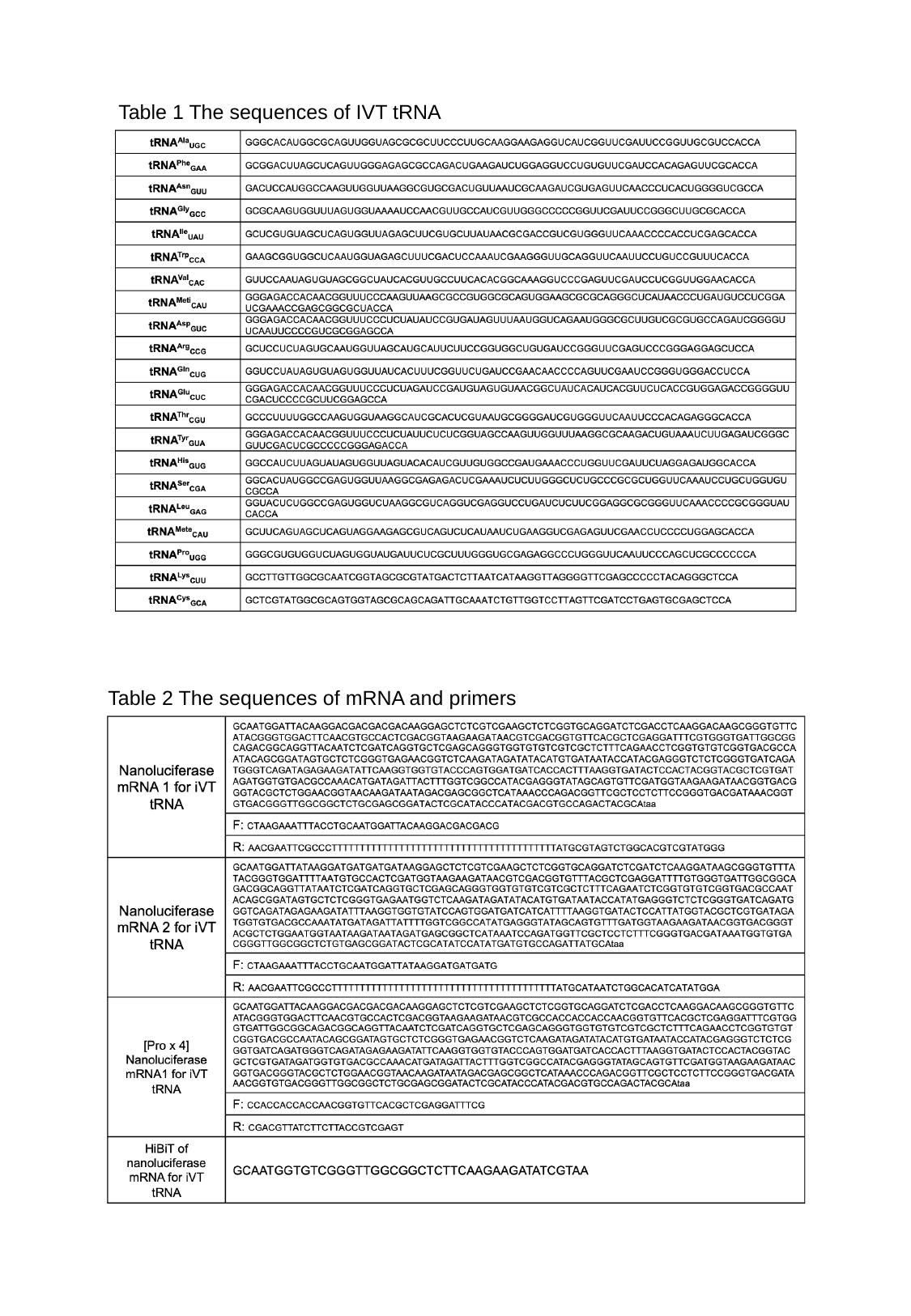

Table 1 The sequences of IVT tRNA
Table 2 The sequences of mRNA and primers
